# De novo lipogenesis fuels adipocyte autophagosome membrane dynamics

**DOI:** 10.1101/2022.05.25.493413

**Authors:** Leslie A. Rowland, Adilson Guilherme, Felipe Henriques, Chloe DiMarzio, Nicole Wetoska, Mark Kelly, Keith Reddig, Gregory Hendricks, Meixia Pan, Xianlin Han, Olga R. Ilkayeva, Christopher B. Newgard, Michael P. Czech

**Affiliations:** Program in Molecular Medicine, University of Massachusetts Chan Medical School, Worcester, MA, 01605, USA; Department of Radiology, University of Massachusetts Chan Medical School, Worcester, MA, 01605, USA; Barshop Institute for Longevity and Aging Studies, University of Texas Health Science Center at San Antonio, San Antonio, TX, 78229, USA; Department of Medicine, University of Texas Health Science Center at San Antonio, San Antonio, TX, 78229, USA; Duke Molecular Physiology Institute, Department of Medicine, Division of Endocrinology, Metabolism and Nutrition, Duke University School of Medicine, Durham, NC, 27710, USA

## Abstract

Autophagy is a homeostatic degradative process for cell components that enables stress resilience and can determine cellular fate and function. However, lipid sources for the rapid membrane expansions of autophagosomes, the workhorses of autophagy, are poorly understood. Here, we identify de novo lipogenesis (DNL) as a critical source of fatty acids (FA) to fuel autophagosome dynamics in adipocytes. Adipocyte fatty acid synthase (Fasn) deficiency markedly impairs autophagy, evident by autophagosome accumulation, and severely compromises degradation of the autophagic substrate p62. Autophagy dependence on FA produced by Fasn is not fully alleviated by exogenous FA in cultured adipocytes even though lipid droplet size is restored. Imaging studies reveal that Fasn colocalizes with nascent autophagosomes, while loss of Fasn decreases certain membrane phosphoinositides known to be required for autophagosome assembly. Together, our studies highlight a newly appreciated function for adipocyte DNL in autophagosome membrane formation and provide evidence that localized FA synthesis contributes to autophagosome dynamics.

## Introduction

Adipose tissues are major regulators of whole-body metabolism, and disruptions to adipose tissue function can drastically alter overall health (Lim et al., 2021). White adipose tissue (WAT) sequesters energy in the form of triglycerides (TGs), which are derived from two main sources of lipid: exogenous circulating fatty acids and endogenously synthesized fatty acids arising from de novo lipogenesis (DNL). Through DNL, fatty acids are synthesized from acetyl-CoA derived from carbohydrates, amino acids, and other sources. Specifically, acetyl-CoA is converted to malonyl-CoA by acetyl-CoA carboxylase (ACC), and these substrates combine to form palmitate through the activities of fatty acid synthase (Fasn) (Wallace and Metallo, 2020). Palmitate is in turn the precursor for synthesis of many other fatty acid types. In adipose tissue, DNL is thought to serve mostly for storing excess energy from carbohydrates, amino acids and other carbon sources as fatty acids, which become esterified and sequestered into TGs (Song et al., 2018). This TG storage function of WAT is hypothesized to enhance whole body metabolic health by enhancing glucose disposal and sequestering toxic lipids away from metabolic tissues such as liver and skeletal muscle (Czech, 2017; Czech et al., 2013; Krahmer et al., 2013; Lotta et al., 2017).

Although the above pathways are well established, adipocyte DNL surprisingly accounts for <10% of adipose TG content; instead, most adipocyte TGs are derived from circulating TGs found in lipoproteins (Czech et al., 2013; Strawford et al., 2004). Supporting the notion that DNL contributes minimally to TG in fat stores, loss of adipocyte Fasn, the rate-limiting enzyme of DNL, has little effect on total lipid accumulation in adipose tissues in mice under normal feeding conditions (Lodhi et al., 2012). Instead, adipocyte specific Fasn knockout mice exhibit browning of white adipose tissue and improved glucose homeostasis (Guilherme et al., 2017; Lodhi et al., 2012). These data raise the idea that DNL and Fasn may be playing important roles in adipocytes beyond merely being a source of fatty acids for storage in lipid droplets. In fact, in a myriad of cell types, Fasn is often necessary for cell survival or proper cellular function. Some of these functions include maintenance of native sarcoplasmic reticulum membrane composition in muscle and plasma membrane cholesterol in macrophages and retina, Schwann cell myelination, and production of ligands for PPAR transcription factors (Funai et al., 2013; Lodhi et al., 2012; Montani et al., 2018; Rajagopal et al., 2018; Wei et al., 2016). The main cellular role of Fasn in adipocytes, however, has remained a mystery.

The studies described here identify a critical role for Fasn in adipocyte autophagy. Autophagy is a degradative process in which cellular components are broken down and recycled by lysosomes (Lamb et al., 2013). Autophagy can promote survival under stress conditions, such as a means of recycling nutrients in response to starvation, but also serves homeostatic roles in the clearance of damaged or unneeded cellular components (Lamb et al., 2013; Melia et al., 2020). During autophagy, cellular contents destined for degradation are encompassed and sequestered by autophagosomes, double-membrane vesicles that ultimately fuse with lysosomes. Autophagosomes form on-demand, and it’s estimated their growth requires an astonishing ∼4,000 phospholipids per second, but the source of these membrane lipids has largely remained elusive (Chang et al., 2021). Our results presented here demonstrate that fatty acids produced by Fasn are a critical source of lipids for these remarkable autophagosome dynamics. This conclusion is consistent with studies in both yeast and mammalian cells that have suggested a link between de novo lipid synthesis and the growing phagophore (Andrejeva et al., 2020; Gross et al., 2019; Nishimura et al., 2017; Schutter et al., 2020). Indeed, we show that adipocytes require localized fatty acid synthesis for efficient autophagy, as exogenous fatty acid supplementation does not fully rescue the defects observed in Fasn-deficient cells. These new data demonstrate a key role of adipocyte DNL in autophagosome membrane expansion, as opposed to its relatively minor contribution to adipocyte TG storage.

## Results

### Fasn deficient adipocytes display impaired autophagy

In contrast to adipocytes in vivo, cultured adipocytes that are differentiated from progenitor cells (preadipocytes) in vitro rely almost exclusively on DNL for TG synthesis and lipid droplet formation due to the low fatty acid content of cell culture media. Therefore, we reasoned that investigating Fasn function might be most productively achieved in cultured adipocytes. Although Fasn is required for adipocyte differentiation, Fasn deficient adipocytes can be obtained from preadipocytes derived from Fasn^Fl/Fl^ mice harboring adiponectin promoter-driven Cre recombinase. In such cells, Fasn deletion is delayed until after onset of adipocyte differentiation, enabling adipocyte formation. Indeed, preadipocytes from Fasn^Fl/Fl^ and Fasn^Fl/Fl^; Adiponectin-Cre+ (cAdFasnKO) mice differentiate competently into adipocytes (Figure 1A). However, whereas cAdFasnKO adipocytes initially accumulate lipid droplets in a manner similar to Fasn-expressing control cells, they lose most of their lipids by Day 7 post-differentiation (Figure 1A). Interestingly, this loss of lipids does not appear to be accompanied by a de-differentiation of the cells, as they maintain their rounded adipocyte morphology and express genes known to be abundant in adipocytes (Figure 1B).

**1.**
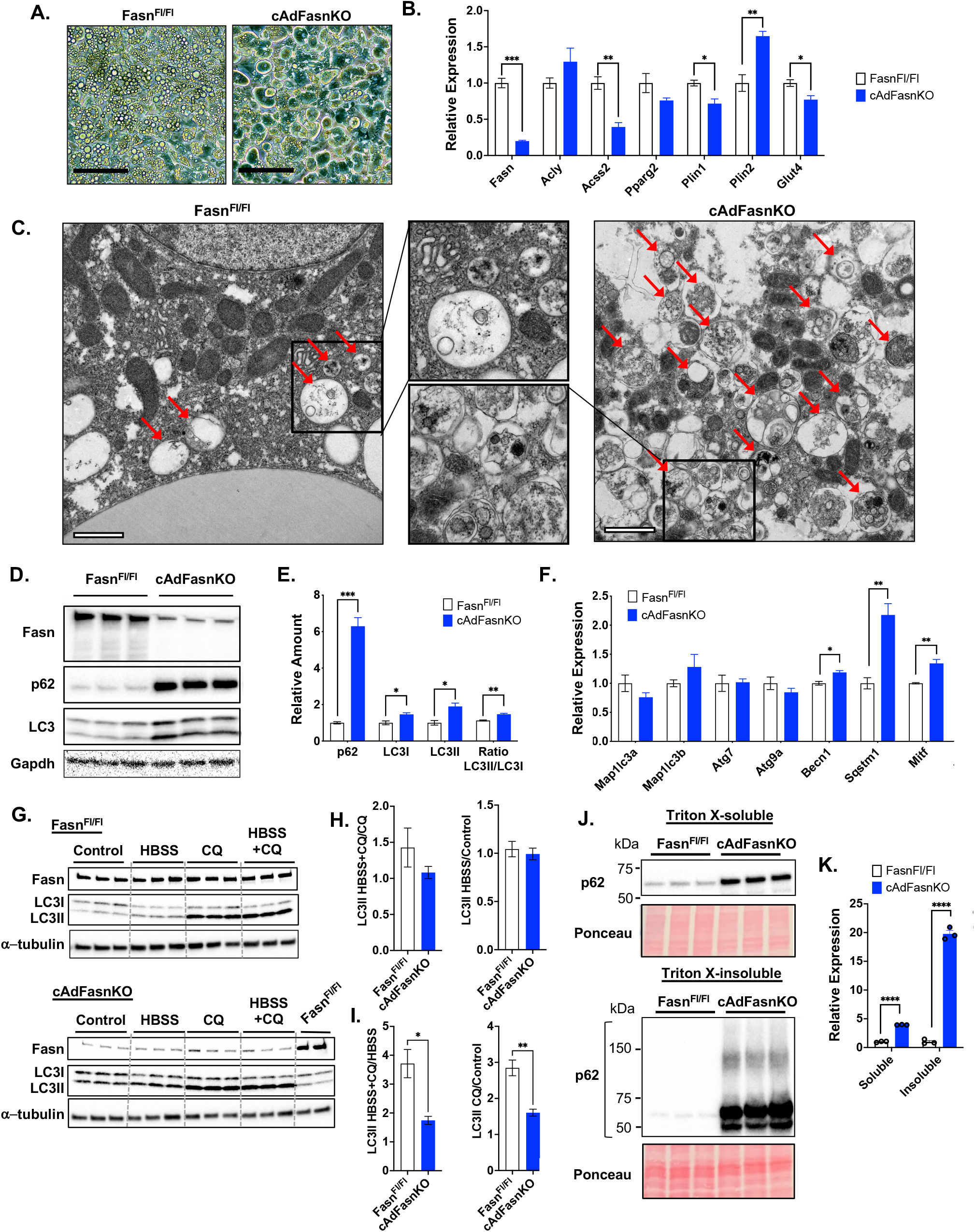
Impaired autophagy in adipocytes deficient in Fasn. A) Light microscopy of in vitro differentiated Fasn^Fl/Fl^ and cAdFasnKO primary adipocytes. Scale bar = 50um. B) qPCR of DNL genes and adipocyte markers C) Electron micrographs of same adipocytes as in A. Arrows indicate autophagic vesicles. Scale bar = 1um. D) Western blot and E) quantification of autophagic markers of Fasn^Fl/Fl^ and cAdFasnKO adipocytes. Proteins normalized to Gapdh. F) qPCR of autophagy-related genes in Fasn^Fl/Fl^ and cAdFasnKO primary adipocytes. *Map1lc3a*=LC3A, *Map1lc3b*=LC3B, *Sqstm1*=p62. G) Western blots of LC3II turnover assay in Fasn^Fl/Fl^ and cAdFasnKO adipocytes. Cells were starved (in HBSS) and/or treated with CQ for 4hrs. HBSS = Hanks’ balanced salt solution, CQ = 50µM chloroquine. H) Synthesis ratio of LC3II calculated as the ratio of LC3II with HBSS to the no HBSS condition. The ratio was calculated with and without autophagy inhibition with CQ as indicated. I) Degradation ratio of LC3II calculated as the ratio of LC3II in the presence of CQ to no CQ treatment. The ratio was calculated with and without autophagy induction with HBSS as indicated. J) Western blots for p62 of Triton X-100-soluble and -insoluble protein fractions. Ponceau S staining provided for loading control. K) Quantification of blots in J. All data are means +/- SE. t tests: *<0.05, **<0.01, ***<0.001, ****<0.0001

Imaging of Fasn-deficient adipocytes by electron microscopy reveals a striking abundance of autophagic vesicles, characterized by a double membrane surrounding cytoplasmic content/organelles in various states of degradation (Figure 1C). Accumulation of autophagic vesicles correlated as expected with a clear increase in levels of the autophagosome marker LC3 (Figure 1D and 1E). We also assessed levels of p62, a mediator of selective autophagy and an autophagy substrate. Levels of this protein were strongly increased (∼6-fold) in cAdFasnKO adipocytes (Figure 1D and 1E), which could not be fully attributed to a much smaller increase (∼2-fold) in levels of the p62/*Sqstm1* transcript (Figure 1F), suggesting a potential stabilization of p62 or a decrease in its degradation. Immunofluorescence labeling also showed strong increases in LC3 and p62 protein in cAdFasnKO adipocytes (Figure S1A). Gene expression analysis showed that *Becn1* and *Mitf* transcripts were slightly elevated in KO adipocytes, whereas expression of other autophagy-related genes was unaffected (Figure 1F).

After proteolytic processing from a common precursor (proLC3), the LC3 protein exists in two forms in the cell: LC3I and LC3II. LC3II differs from LC3I in that upon autophagy activation, the LC3I protein becomes conjugated to phosphatidylethanolamine, forming LC3II. LC3II is then anchored into the autophagosome membrane and thus serves as a reliable marker of both growing and completed autophagosomes. Because an increase in LC3II could indicate an increase in autophagosome formation but also a decrease in autophagosome degradation, measuring changes in LC3II in response to stimuli that activate or block autophagy is necessary to adequately assess autophagic flux (Klionsky et al., 2021). The increases in LC3 and p62 proteins in cAdFasnKO adipocytes suggesting altered autophagy flux prompted us to investigate further with an LC3II turnover assay. Adipocytes were treated with HBSS to activate autophagy and chloroquine (CQ) to block autophagic degradation. Shown in Figure 1G and quantified in Figure S1B, activation of autophagy with HBSS had no effect on LC3II levels in either control or KO adipocytes in the absence of CQ. Blockade of autophagic degradation by CQ, however, increased LC3II in WT and KO adipocytes (Figure 1G and S1A). This large increase in LC3II accumulation with CQ but not with HBSS indicates these adipocytes likely already have a high rate of autophagic flux under basal conditions. Since LC3II levels differ between Fasn^Fl/Fl^ and cAdFasnKO adipocytes under basal conditions (Figure 1D and 1G), it is necessary to calculate the ratios of LC3II in the presence of HBSS and CQ to accurately determine flux (Klionsky et al., 2021). The LC3II synthesis ratio, a measure of autophagosome formation rate, is not significantly reduced by FasnKO (Figure 1H). However, calculation of the LC3II degradation ratio, a measure of autophagic degradation, showed a significant reduction by FasnKO (Figure 1I). These data indicate that autophagic flux is impaired in cAdFasnKO adipocytes due to reductions in autophagic degradation.

Since p62 is an autophagy substrate, p62 degradation can be another useful way to assess autophagy flux in cells. Under conditions in which autophagy is inhibited, p62 degradation is impaired, and accumulated p62 condenses to form detergent-insoluble aggregates (Klionsky et al., 2021). We assessed the presence of these insoluble aggregates by fractionating adipocyte protein lysates into Triton X-100-soluble and Triton X-100 insoluble forms. In the soluble fraction of Fasn KO cells, p62 was significantly increased similar to the data in Figure 1D, in concert with massive accumulation of p62 in the insoluble fraction (Figure 1J and 1K). These data further confirm autophagic degradation is strongly impaired in cAdFasnKO adipocytes.

### Protein malonylation is increased in cAdFasnKO adipocytes

While loss of Fasn eliminates the production of palmitate via DNL, it may also affect levels of earlier pathway intermediates, such as acetyl-CoA and malonyl-CoA (Bruning et al., 2018; Chakravarthy et al., 2005) (Figure 2A). The role of malonyl-CoA in autophagy has not been determined, whereas acetyl-CoA is a well-established negative regulator of autophagy. Reductions in cytosolic acetyl-CoA activate, while increases in acetyl-CoA inhibit autophagy (Marino et al., 2014). For these reasons, it was important to establish whether the impairment of autophagy in cAdFasnKO adipocytes might be linked to reduced palmitate production or altered levels of acetyl-CoA or malonyl-CoA (Figure 2A).

**2.**
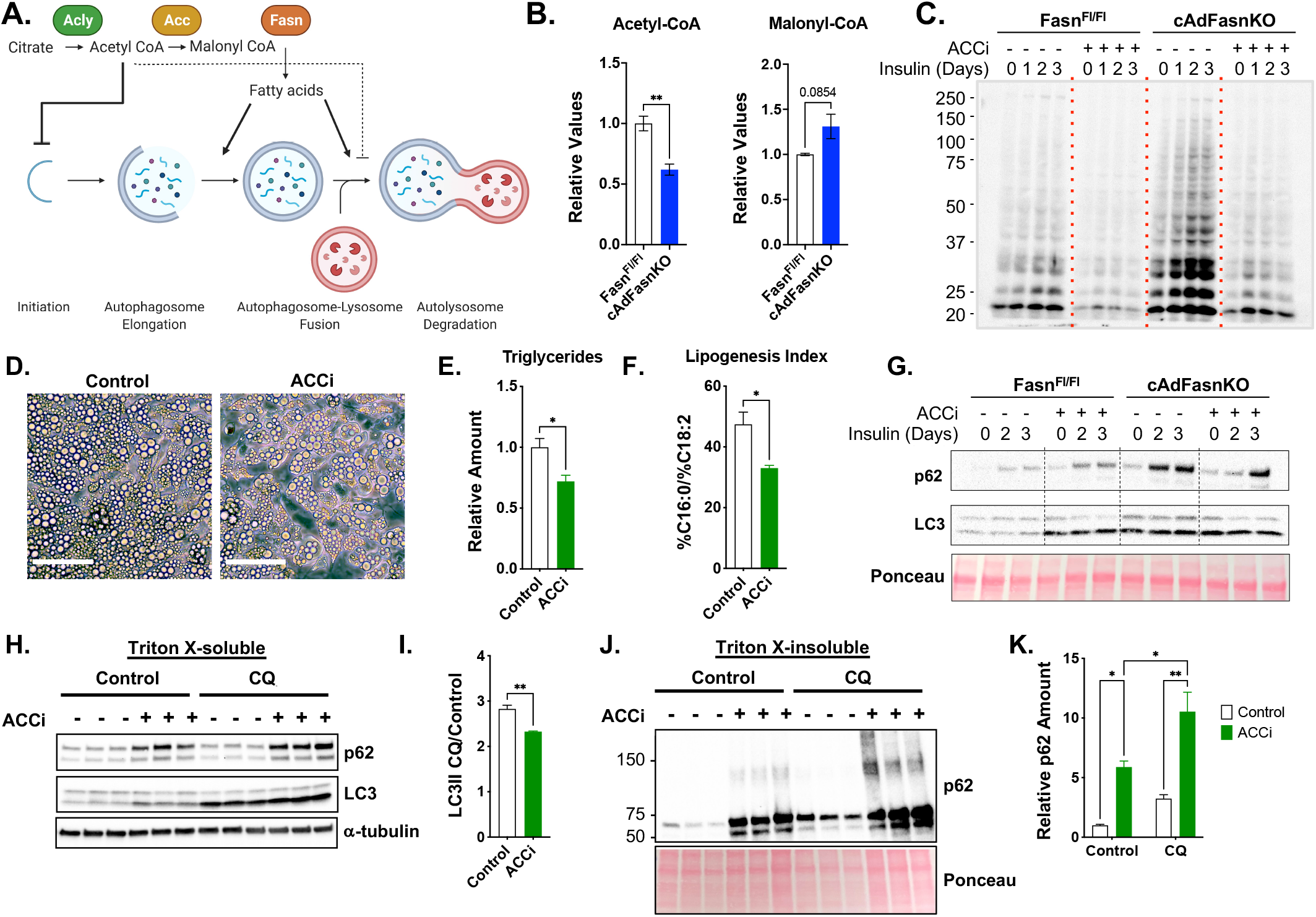
DNL metabolites acetyl-CoA and malonyl-CoA do not mediate the impaired autophagy in cultured FasnKO adipocytes. A) Diagram of de novo lipogenesis and its proposed role(s) in autophagy. B) Relative acetyl-CoA and malonyl-CoA levels in Fasn^Fl/Fl^ and cAdFasnKO adipocytes determined by LC-MS/MS. t test: **<0.01 C) Western blot of malonylated proteins in Fasn^Fl/Fl^ and cAdFasnKO adipocytes in the presence of 15µM ND-630 (ACC inhibitor, ACCi) for 72 hours and 1µM insulin for the indicated time points. D) Light microscopy of in vitro differentiated adipocytes treated with and without 10µM ACCi for 48 hours. Scale bar = 50µm E) Relative triglyceride content and F) Lipogenesis index calculated as the ratio of the percentage of C16:0 to the percentage of C18:2 in triglycerides in control and ACCi-treated adipocytes. t test: *<0.05 G) Western blot of soluble p62 and LC3 in Fasn^Fl/Fl^ and cAdFasnKO adipocytes treated with ACCi and insulin as shown in C. H) Western blot for p62 and LC3 of Triton X-100-soluble protein fractions from adipocytes treated with 10µM ACCi for 48 hours with and without 4-hour treatment with 50µM CQ I) LC3II degradation ratio calculated from Figure 2H. t test: **<0.01, J) Western blot for p62 of Triton X-100-insoluble protein fractions from adipocytes treated with 10µM ACCi for 48 hours with and without 4-hour treatment with 50µM CQ K) Quantification of insoluble p62 from 2J. Ponceau provided as loading control. 2-way ANOVA with Tukey post hoc: *<0.05, **<0.01. All data are means +/- SE.

To determine whether acetyl-CoA or malonyl-CoA levels were affected by Fasn deletion, LC-MS/MS was performed on cell extracts. Surprisingly, total acetyl-CoA levels were significantly reduced, while malonyl-CoA was slightly but not significantly elevated in cAdFasnKO adipocytes (Figure 2B). Immunoblot analysis of total malonylated proteins demonstrated a dramatic increase in cAdFasnKO adipocytes (Figure 2C). This discrepancy between malonyl-CoA levels and malonylated proteins may suggest that protein malonylation serves as a sink for excess malonyl-CoA. In contrast, Western blotting for total acetylated proteins showed minimal effects of Fasn KO in adipocytes (data not shown). These data indicate that acetyl-CoA is unlikely to play a significant role in the autophagy inhibition observed in cAdFasnKO adipocytes.

### Protein malonylation in cAdFasnKO adipocytes does not mediate autophagy inhibition

To determine whether protein malonylation or malonyl-CoA impairs autophagy, we used ND-630, a pharmacological inhibitor of ACC1/2, the enzyme responsible for malonyl-CoA production immediately upstream of Fasn (Figure 2A). Inhibition of ACC in fully differentiated cultured adipocytes caused a slight diminution of lipid droplets as observed by light microscopy (Figure 2D), accompanied by a reduction in cellular triglycerides (Figure 2E). Quantification of the lipogenesis index, the ratio of C16:0 to C18:2 in triglycerides and a measure of DNL (Chong et al., 2008; Hudgins et al., 1996), indicated that DNL was effectively inhibited by ND-630 treatment (Figure 2F). ND-630 treatment also successfully reduced protein malonylation in control cells and returned protein malonylation levels to control levels in cAdFasnKO adipocytes (Figure 2C). Together, these data demonstrate that pharmacological inhibition of ACC in fully differentiated adipocytes effectively reduces protein malonylation and DNL.

Western blotting for autophagy markers LC3 and p62 showed that ACC inhibition alone is sufficient to increase LC3II and p62 content (Figure S2A and S2B), while ACC inhibition in cAdFasnKO adipocytes does not restore LC3II or p62 to control levels, indicating increased protein malonylation does not contribute to autophagy impairment (Figure 2G). Since we noticed an effect of ND-630 alone on autophagy markers, we measured autophagy flux in the ND-630-treated adipocytes in the presence of CQ. Similar to the findings with cAdFasnKO adipocytes, ACC inhibition significantly reduces autophagy flux measured by the LC3II degradation ratio (Figure 2H and 2I). ACC inhibition also significantly increases both soluble and insoluble p62 levels (Figure 2H and 2J, quantified in Figure S2A and 2K, respectively). Combined, these data show that like cAdFasnKO, ACC inhibition in adipocytes significantly impairs autophagic degradation, providing independent evidence that inhibition of fatty acid synthesis is sufficient to impair autophagy.

### Fatty acids produced by Fasn are essential for autophagy

If fatty acid synthesis via Fasn is required for autophagy, we hypothesized that the provision of exogenous fatty acids would restore autophagy function in cAdFasnKO adipocytes. Supplementation of cAdFasnKO adipocytes with a mixture of palmitate and oleate indeed restored soluble p62 levels, though insoluble p62 levels were only partially rescued (Figure 3A and 3B). Likewise, LC3II was not restored by the addition of fatty acids to cAdFasnKO adipocytes (Figure 3A and 3C). Similar results were obtained when fatty acids were provided for a longer period and throughout the differentiation process, where lipid droplets were plentiful in KO adipocytes (Figure S1A-D). Thus, the autophagy impairment in cAdFasnKO adipocytes could not be ascribed simply to a lack of adipocyte lipid content. Importantly, *p62/Sqstm1* mRNA levels were not reduced by fatty acids (Figure 3D), though other autophagy gene expression changes as a result of cAdFasnKO were restored by fatty acids (Figure 3E). Immunofluorescent labeling similarly showed a partial reduction of p62 levels in cAdFasnKO adipocytes by fatty acids (Figure 3F). These data suggest that the partial rescue of p62 protein levels was a direct consequence of fatty acids restoring p62 degradation and thus autophagy flux in FasnKO adipocytes.

**3.**
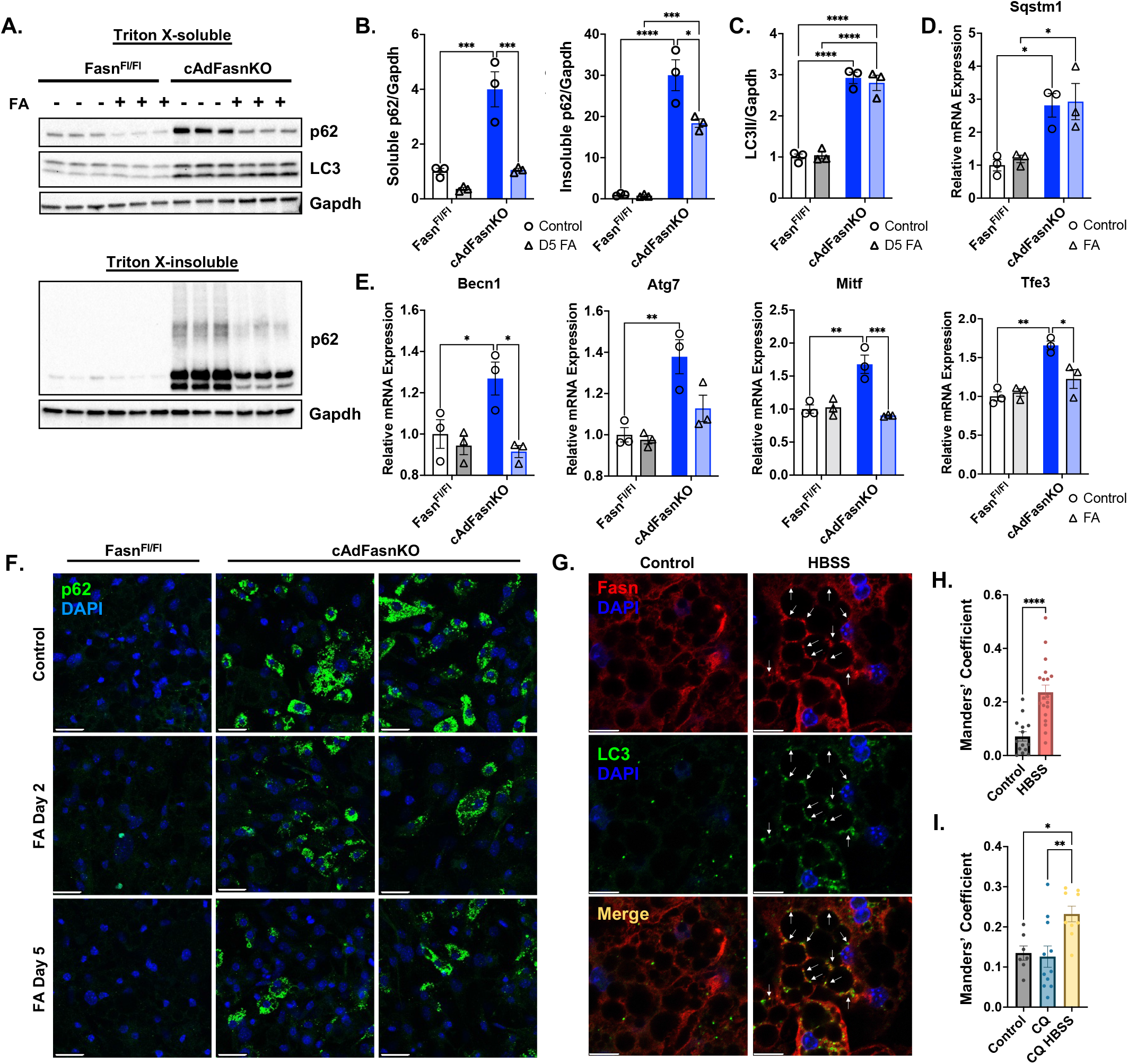
Fatty acids produced by Fasn are essential for autophagy. Fasn^Fl/Fl^ and cAdFasnKO adipocytes were supplemented with 200uM each of palmitate and oleate for 48 hours after they were fully differentiated (day 5). A) Western blots of Triton X-100-soluble and -insoluble protein fractions. B) Quantification of soluble and insoluble p62 in Figure 3A normalized to Gapdh. C) Quantification of LC3II normalized to Gapdh from Figure 3A. D) qPCR of *Sqstm1/*p62 and E) autophagy-related genes in adipocytes supplemented with fatty acids (FAs) throughout differentiation, beginning on day 0. All data are means +/- SE. All datasets analyzed with two-way ANOVA with Tukey post hoc: *<0.05, **<0.01, ***<0.001, ****<0.0001. F) Immunofluorescence of Fasn^Fl/Fl^ and cAdFasnKO adipocytes supplemented with FAs beginning on Day 2 or Day 5 of differentiation and labeled with p62. Scale bar = 50um. G) Immunofluorescence imaging of wild-type adipocytes starved for 2 hours in HBSS and double-labeled for LC3B and Fasn. Scale bar = 10um H) Quantification of colocalization of LC3B and Fasn in Figure 3F as measured by Manders’ coefficient = fraction of LC3B colocalized with Fasn. Data are means +/- SE. t test: ****<0.0001. I) Manders’ coefficient for adipocytes treated with 30uM CQ for 24 hours (CQ condition) or starved in HBSS with 50uM CQ for 5 hours (CQ HBSS condition). Data are means +/- SE. One-way ANOVA with Sidak post hoc. *<0.05, **<0.01.

Because exogenous fatty acids and lipid droplet restoration were not sufficient to fully rescue autophagic flux in knockout adipocytes, we hypothesized that local fatty acid synthesis is required for proper autophagosome dynamics. To test this, we performed immunofluorescence colocalization analyses to determine if Fasn colocalized with the autophagosome marker, LC3B, shown in Figure 3G. As expected, LC3B puncta formation was increased by starvation in HBSS (Figure 3F). Fasn immunolabeling showed diffuse cytoplasmic staining but also appeared in punctate structures, which were increased by autophagy-activating conditions; i.e. starvation. Interestingly, when autophagy was activated by starvation, the amount of LC3B colocalized with Fasn significantly increased (Figure 3H). Importantly, increasing LC3B puncta formation by autophagy inhibition (CQ) did not increase Fasn/LC3B colocalization; whereas, autophagy activation under these conditions did increase colocalization (Figure 3I). These data indicate Fasn localizes with nascent autophagosomes and likely contributes to the growing autophagosome membrane. Together, these data are consistent with the hypothesis that endogenous fatty acid synthesis directly supplies lipids to autophagosome membranes and is required for effective autophagic flux.

### Impaired autophagy in adipocytes of cAdFasnKO mice

We next asked whether the autophagy defect in Fasn deficient adipocytes could be observed in vivo. For these experiments, we used tamoxifen-inducible, adipocyte specific FasnKO (iAdFasnKO) mice, as well as the constitutively active Adiponectin-Cre (cAdFasnKO) mice that were used for the previous studies on cultured adipocytes. In subcutaneous white adipose tissue of iAdFasnKO mice, we observed increased p62 and LC3II (Figure 4A), consistent with our experiments performed in vitro. Also observed was the characteristic beiging of iAdFasnKO mice evident by increased Ucp1 expression. Similar results were found after mice were fasted, an autophagy-activating condition (Figure 4A). These changes in autophagic markers occurred specifically in adipocytes, shown by fractionation of adipose tissue (Figure 4B) and immunohistochemical staining of p62 in WAT (Figure 4C). Increased p62 protein was observed in all adipose tissues but was not due to increased *Sqstm1* transcription (Figure 4D and 4E, respectively). An ex vivo autophagy flux assay assessing LC3II and p62 changes in the presence of CQ showed significantly increased LC3II and p62 in both basal and CQ-treated conditions (Figure 4F and 4G). Calculation of the degradation ratio for LC3II indicated a tendency for a reduced autophagosome degradation in KO explants (Figure 4H), although there was a significant reduction in p62 degradation (Figure 4H). These results suggest an impairment in autophagosome degradation in iAdFasnKO adipose tissue in vivo, consistent with our findings in cultured cAdFasnKO adipocytes in vitro.

**4.**
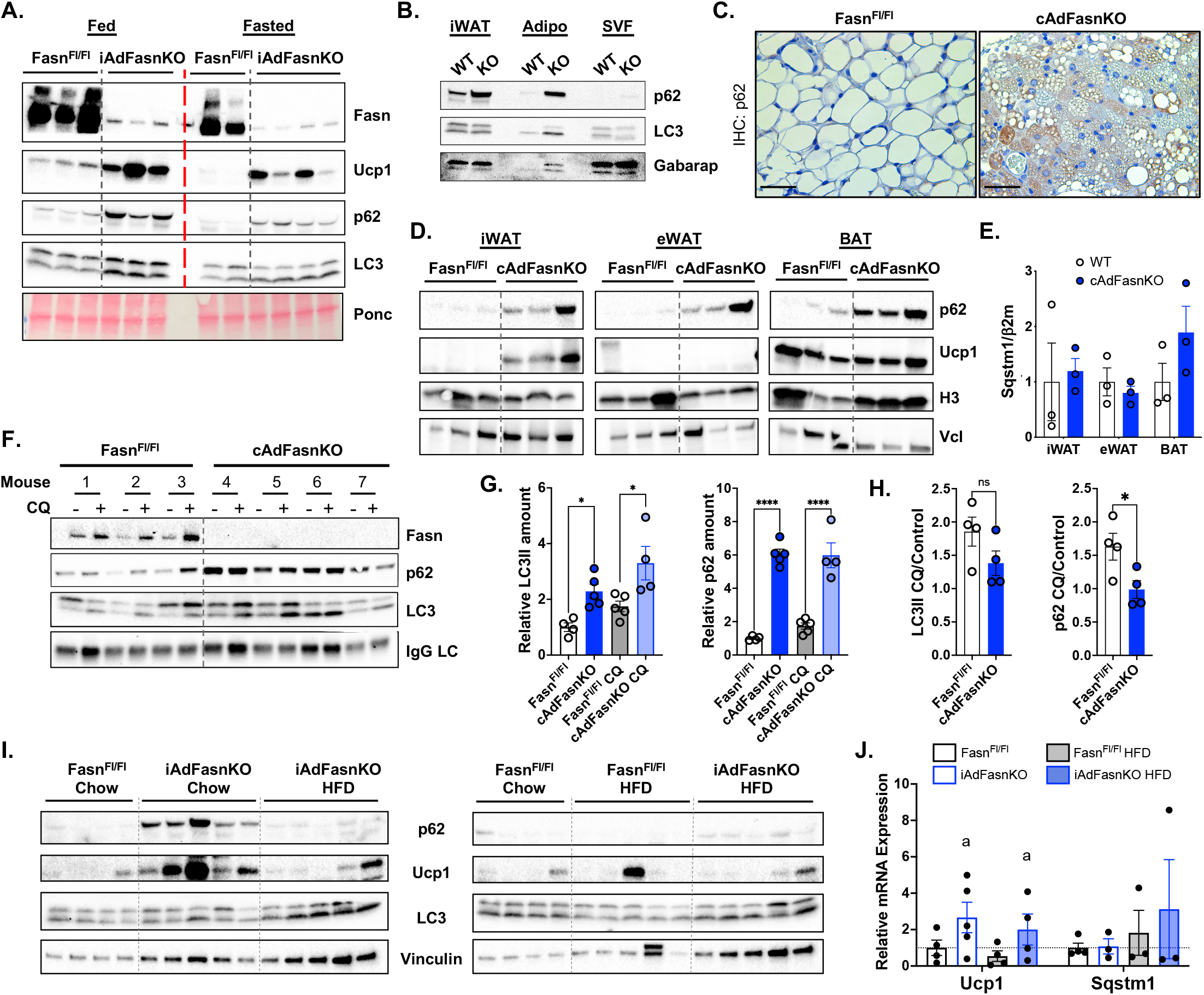
Adipocyte Fasn deficiency impairs autophagy and induces accumulation of p62 protein in vivo. A. Western blot of subcutaneous white adipose tissue (WAT) from Fasn^Fl/Fl^ and inducible AdFasnKO (iAdFasnKO) mice under fed and 24-hour fasted conditions. Ponceau (Ponc) provided for loading control. B) Western blot of pooled subcutaneous WAT (iWAT) from Fasn^Fl/Fl^ (WT) and iAdFasnKO (KO) mice separated into adipocyte (Adipo) and stromal vascular fractions (SVF). C) Immunohistochemical staining of p62 in iWAT of Fasn^Fl/Fl^ and iAdFasnKO mice; scale bar = 25um. D) p62 Western blot of adipose tissue depots from Fasn^Fl/Fl^ and cAdFasnKO mice. Histone H3 (H3) and vinculin (Vcl) provided as loading controls. eWAT = epididymal adipose tissue, BAT = brown adipose tissue. E) qPCR comparing p62 (*Sqstm1*) mRNA expression across adipose tissue depots. F) Ex vivo autophagy flux assay in subcutaneous WAT explants harvested from Fasn^Fl/Fl^ and cAdFasnKO mice. WAT explants were cultured in serum-free DMEM/F12 media with or without 50uM CQ for 2 hours. Western blot results are shown. G) Quantification of LC3II and p62 in western blot in Figure 4F. Proteins were normalized to mouse IgG light chain levels. 2-way ANOVA with repeated measures and Sidak post hoc: *<0.05, ****<0.0001. H) Degradation ratios of LC3II and p62 were determined by calculating the change in protein amount in the CQ condition divided by the control condition. t tests: *<0.05 I) Fasn^Fl/Fl^ and iAdFasnKO mice fed a chow diet or high fat diet. Western blot of subcutaneous WAT. J. qPCR of *Ucp1* and *Sqstm1* (p62) in samples from 4I. Two-way ANOVA: a = main effect of FasnKO. All data are means +/- SE.

Based on the partial restoration of autophagy flux by fatty acid supplementation in vitro, mice were fed a high fat diet (HFD) rich in saturated fatty acids to determine if this would restore autophagy flux in vivo. Shown in Figure 4I, HFD feeding reduced p62 protein levels and interestingly also blunted Ucp1 protein levels in iAdFasnKO iWAT. Again, these p62/Sqstm1 changes were not due to transcriptional effects (Figure 4J). Changes in LC3 lipidation were not as clear in this cohort of chow-fed iAdFasnKO mice. We hypothesize these variations are due to the cellular heterogeneity of the adipose tissue in vivo as we see a significant amount of LC3 in the SVF fraction (Figure 4B). These data in mice reinforce our findings with cAdFasnKO adipocytes in culture indicating Fasn is required for proper autophagy flux in adipocytes, and exogenous fatty acids can only partially restore autophagic function.

### Deletion of Fasn alters the cellular lipidome in vitro and in vivo

Given that deletion of Fasn appears to interfere with the degradation of autophagic vesicles but not their formation, Fasn may function to ensure proper membrane lipid composition for efficient autophagy dynamics. We therefore performed lipidomics analysis on Fasn deficient adipocytes to investigate the effect of Fasn on cellular lipid composition. In vitro, loss of adipocyte Fasn led to significant remodeling of phospholipid composition. Phosphatidylinositol, palmitic acid, cardiolipin, and all measured lysophospholipid species were reduced, while phosphatidylethanolamine, phosphatidylglycerol, and ceramides were increased (Figure 5A). In vivo, however, only sphingomyelin was significantly reduced in adipose tissue (Figure 5B).

**5.**
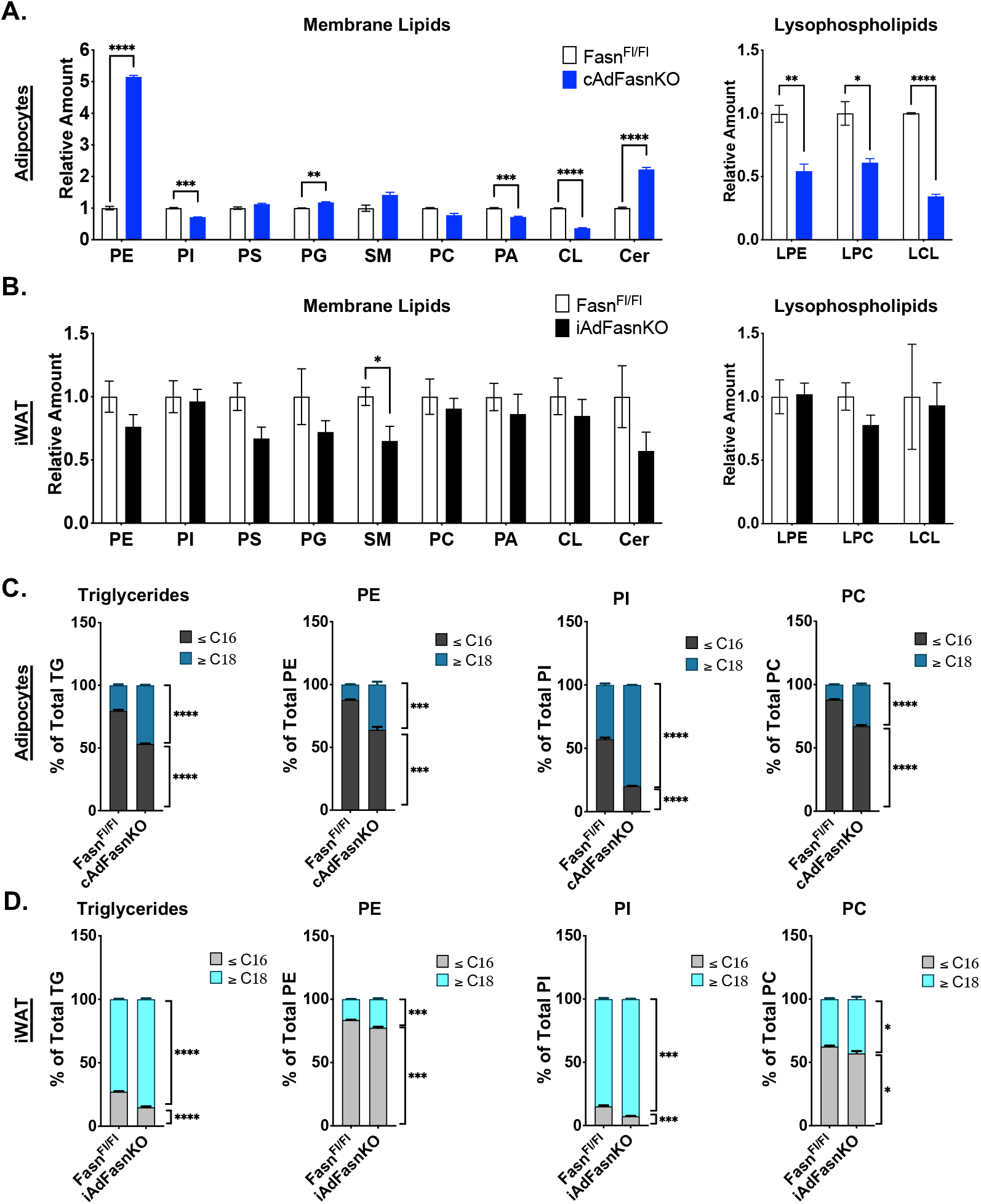
Deletion of Fasn alters the cellular lipidome in vitro and in vivo. Lipidomics analyses from in vitro differentiated Fasn^Fl/Fl^ and cAdFasnKO primary adipocytes and subcutaneous WAT from Fasn^Fl/Fl^ and iAdFasnKO mice. A) Relative amounts of different lipid classes from in vitro differentiated adipocytes and B) in vivo iWAT. t tests: *<0.05, **<0.01, ***<0.001, ****<0.0001. C,D) Fatty acyl chain composition of different lipid classes in vitro (C) and in vivo (D). Percent of indicated lipid containing fatty acid chains </= 16 carbons vs. lipids with fatty acid chains >/= 18 carbons. All data are means +/- SE. t tests: *<0.05, **<0.01, ***<0.001, ****<0.0001. PE, phosphatidylethanolamine; PI, phosphatidylinositol; PS, phosphatidylserine; PG, phosphatidylglycerol; SM, sphingomyelin; PC, phosphatidylcholine; PA, phosphatidic acid; CL, cardiolipin; Cer, ceramide; LPE, lysophosphatidylethanolamine, LPC, lysophosphatidylcholine; LCL, lysocardiolipin.

In mice, the effect of FasnKO on membrane lipids is likely obscured by a constant delivery of fatty acids from the circulation. This prompted us to examine the fatty acyl composition of the lipidome to determine if Fasn is contributing specific fatty acids to each lipid species. We found that Fasn deficiency consistently caused a shift in fatty acyl composition toward longer chain (>18C) fatty acids in various lipid classes both in vitro (Figure 5C) and in vivo (Figure 5D), reflecting a reduction in palmitate (C16:0) synthesis. We also found that loss of Fasn results in shifts in double bond composition in vitro and in vivo, where FasnKO adipocytes contain fatty acyl chains with greater numbers of double bonds, which arise from essential fatty acids taken up from culture media or dietary intake (Figure S2A and S2B).

Phosphatidylinositols are critical for multiple steps of the autophagy pathway, from autophagosome formation to fusion with the lysosome (Zhao et al., 2021; Zhao and Zhang, 2019). Investigation of the specific phosphatidylinositol species altered by Fasn deficiency in vitro and in vivo showed that PI(16:0-18:2), PI(16:1-20:4), and PI(16:0-20:4) were significantly reduced and PI(18:0-20:4) was significantly increased (Figure S2C) suggesting these particular PI species may play specific roles in autophagy. Collectively, these data show that loss of Fasn can significantly alter the cellular lipidome and can alter specific phospholipids both in vitro and in vivo.

## Discussion

The major finding of this study is the identification of de novo lipogenesis as an essential source of lipids for autophagy in adipocytes. It has previously been appreciated that autophagosome synthesis is an extremely lipid-demanding process, yet the precise source(s) of these lipids has not been fully elucidated (Chang et al., 2021; Melia et al., 2020). Data presented here demonstrate that endogenous de novo fatty acid synthesis via Fasn funnels fatty acids into autophagosomes, and that loss of this pathway disables autophagic degradation. Bioenergetically, our findings are compatible with the concept that it is more favorable to synthesize new membranes than to reuse old membranes, which require extensive processing such as protein removal (Andrejeva et al., 2020). In addition, preferential usage of de novo lipogenesis for membrane lipids rather than depleting preexisting membranes or lipid droplets would appear to confer a benefit to cells, especially during periods of stress. Autophagy serves critical cytoprotective and hormetic roles in virtually all cell types. Thus, our findings answer a fundamental biological question with wide-ranging implications.

Adipose tissue function is paramount to overall health, and the importance of autophagy in adipose tissue function has recently come to light (Cairo and Villarroya, 2020; Ferhat et al., 2019). Excessive autophagy induced by adipocyte-selective deletion of the negative autophagy regulator, Rubicon, results in lipodystrophy, glucose intolerance, and fatty liver. Also, downregulation of Rubicon and the associated increase in autophagy has been linked to the age-dependent decline in adipose tissue and metabolic health (Yamamuro et al., 2020). Conversely, autophagy inhibition has been associated with improvements in metabolic health. Knockout of Atg7, an early autophagy factor, in adipose tissue blocked autophagy and led to reduced body fat, increased insulin sensitivity, and beiging of white adipose tissue (Singh et al., 2009b). In addition, several studies have shown that blocking mitophagy, a specialized form of autophagy, prevents the turnover, or “whitening,” of beige and brown adipose tissue, leading to beneficial metabolic effects (Altshuler-Keylin et al., 2016; Lu et al., 2018; Schlein et al., 2021).

Interestingly, in addition to impaired autophagy, iAdFasnKO and cAdFasnKO mice also exhibit increased white adipose beiging (Figure 4A and 4I) and improved glucose tolerance (Guilherme et al., 2017; Lodhi et al., 2012). One of the more prominent effects of Fasn knockout on adipocytes was the substantial increase in p62 protein (Figure 4A and 4I). In addition to its role in autophagy, p62 serves as a signaling hub that participates in multiple pathways, but most notably, p62 regulates beige and brown adipocyte formation. Knockout of p62 in adipose tissue results in obesity, glucose intolerance, and reduced energy expenditure, effects attributed to reductions in brown and beige adipose tissue thermogenesis (Fischer et al., 2020; Huang et al., 2021; Muller et al., 2013). Mechanistically, p62 directs phospho-ATF2 to promoters of thermogenic genes and also functions as co-activator of PPARγ (Fischer et al., 2020; Huang et al., 2021). It is tempting to speculate that FasnKO-mediated increases in adipose tissue p62 as a result of autophagy dysfunction regulate the beiging and metabolic improvements observed in these mice, but FasnKO in vitro does not upregulate Ucp1 under our experimental conditions. Thus, further investigation is required to determine whether the autophagy impairment in cAdFasnKO mice contributes to development of the beiging phenotype.

A notable finding from these studies was that FasnKO did not impair autophagosome formation, suggested by autophagy flux assays (Figure 1H) and the presence of apparently fully formed autophagosomes in the electron micrographs (Figure 1C). Lipid droplets and preexisting membranes have been shown to contribute to autophagosome membranes, and in the absence of de novo lipogenesis, it’s likely that multiple membrane lipid sources are utilized (Dupont et al., 2014; Melia et al., 2020). Moreover, even though cAdFasnKO adipocytes are largely lipid droplet-depleted, some lipid droplets remain and could serve as lipid sources for autophagosome membranes. Current studies are aimed at identifying the role of Fasn in autophagy and whether autophagosome formation is compromised in non-adipocyte cell types where lipid droplets do not accumulate.

Despite the fact that autophagosomes can form in the absence of Fasn, our data strongly suggest that these autophagosomes are not fully functional, i.e., are not degraded. Autophagic membrane composition must be tightly regulated to ensure proper protein recruitment and membrane dynamics, including autophagosome-lysosome fusion events (Corcelle-Termeau et al., 2016; Koga et al., 2010; Singh et al., 2009a; Zhao et al., 2021; Zhao and Zhang, 2019). Our lipidomics analyses from both cultured adipocytes and adipose tissue showed that Fasn deficiency significantly altered cellular lipid composition, including membrane lipids (Figure 5). In vitro, knockout of Fasn dramatically altered phospholipid levels, including total PI, PA, PG, PE, among others (Figure 5A). Phosphatidylinositol, in particular, plays significant roles in autophagy. Its phosphorylated derivatives, PI3P, PI4P, PI(3,5)P2, serve as recognition and docking sites for autophagy effector proteins on both autophagosome and lysosomal membranes (Zhao et al., 2021; Zhao and Zhang, 2019). Accordingly, reductions in PI synthesis result in impaired autophagy turnover (Nishimura et al., 2017). In addition, alterations to PA content have been shown to affect autophagic flux (Holland et al., 2016). In vivo, however, these same changes to total phospholipids were not evident (Figure 5B), likely due to compensation from uptake of fatty acids from the circulation. Nonetheless, since these lipidomics analyses were performed on whole cells and tissues, we cannot rule out significant alterations to phospholipid profiles at the subcellular level, i.e., in autophagosome membranes, where these changes would be impactful. On the other hand, a common theme between the data obtained in Fasn deficient adipocytes in both cultured cells and in mouse adipose tissue is the significant reduction in C16 and saturated acyl chains across most lipid species, reflecting the reduction in palmitate synthesis (Figure 5C and S4). Alterations to acyl chain length and saturation can have significant effects on membrane fluidity, thickness, and packing density, which can affect fusion with the lysosome (Andrejeva et al., 2020; Holthuis and Menon, 2014; Koga et al., 2010). Together, these data suggest de novo lipogenesis and Fasn equip the autophagosome, and perhaps lysosomal membranes with a precise lipid composition that ensures their proper fusion, trafficking, and degradation.

A prominent finding in this study was that Fasn directly localizes near nascent autophagosomes (Figure 3F and 3G). Supportive of this, our results revealed that fatty acids could not fully restore autophagic function of cultured cAdFasnKO adipocytes (Figure 3A and 3B), and even fully lipid-replete cAdFasnKO adipose tissue exhibited autophagy impairment (Figure 4F-4H). These results indicate that Fasn may be acting locally to supply fatty acids to growing autophagosomes. Our findings are in line with studies in yeast that showed the fatty acyl CoA synthetase, Faa1, directly localized on growing autophagosome membranes (Schutter et al., 2020). The phosphatidylinositol (PI) synthesizing enzyme, PI Synthase (PIS) was also shown to localize to sites of autophagosome biogenesis in mammalian cells (Nishimura et al., 2017). The presence of Fasn and lipid synthesizing enzymes on nascent autophagosomes supports the conclusion that one of the primary functions of Fasn is to directly funnel fatty acids into autophagosome membranes.

In summary, our studies provide the first demonstration of a direct contribution of de novo fatty acid synthesis to autophagosome membrane dynamics in mammalian cells. Our findings are in line with studies implicating the lipid synthesis enzymes Acc1 and Faa1 in yeast and the fatty acid desaturase, Scd1, in mammalian cells in autophagosome membrane synthesis (Gross et al., 2019; Ogasawara et al., 2014; Schutter et al., 2020). These findings could have far-reaching therapeutic implications as inhibitors of de novo lipogenesis enzymes, particularly ACC and Fasn, are currently in development for the treatment of fatty liver and cancers (Batchuluun et al., 2022). Our studies identify an important new function of these enzymes that may impact the usage of these inhibitors. Moreover, dysregulation of autophagy has been implicated in various diseases (Choi et al., 2013). Thus, identification of the de novo lipogenesis pathway as a critical component of autophagy opens new opportunities to modulate autophagy and disease progression.

## Acknowledgements

We thank members of the Czech laboratory for helpful discussions. The electron microscopy in this project was supported by Award Numbers S10OD025113-01 and S10OD021580 from the National Center for Research Resources. Funding was also provided by grants from the National Institute for Diabetes and Digestive and Kidney Diseases (DK116056 and DK030898 to M.P.C. and DK124723 to C.B.N.). The Functional Lipidomics Core at the Barshop Institute for Longevity and Aging Studies is partially supported by center grants from National Institute on Aging (P30 AG013319 and P30 AG044271). The content is solely the responsibility of the authors and does not necessarily represent the official views of the National Center for Research Resources or the National Institutes of Health.

## Author contributions

L.A.R., A.G., and M.P.C. conceived the study and designed the experiments. L.A.R., A.G., C.D., N.W., and M.K. performed the experiments. K.R. and G.H. performed the electron microscopy, M.P. and X.H. performed the lipidomics analyses. O.R.I. and C.B.N. performed the acetyl-CoA and malonyl-CoA measurements and collaborated on interpretations of all the data. L.A.R. and M.P.C. wrote the manuscript, which was reviewed and edited by all co-authors.

## Declaration of interests

The authors declare no competing interests.

## Methods

### Mice

All mice were raised in standard housing conditions and fed a standard chow diet ad libitum under approval by the University of Massachusetts Medical School Institutional Animal Care Use Committee (IACUC). C57Bl/6J (WT) mice were obtained from Jackson Laboratory and bred for primary preadipocyte isolation. Fasn^Flox/Flox^ mice were generated as previously described (cite) and bred with <u>c</u>onstitutively expressed <u>Ad</u>iponectin-Cre (<u>PubMed:21356515</u>; Jax stock #028020) mice to obtain adipocyte-specific <u>Fasn</u> knockout (<u>KO</u>) mice (cAdFasnKO). For inducible knockout of Fasn in adult mice, Fasn^Flox/Flox^ mice were bred with Adiponectin-Cre-ERT2 mice, as previously described. To induce Fasn knockout, Fasn^Flox/Flox^ or Fasn^Flox/Flox^; Adiponectin-Cre-ERT2^+^ mice (iAdFasnKO) were administered via i.p. injection 1mg tamoxifen dissolved in corn oil once per day for 5 days. For high fat diet experiments, mice were fed a 60% kcal from fat diet (Research Diets, D12492i) ad libitum for 4 weeks prior to tamoxifen administration, then remained on the diet for another 12 weeks.

### Primary preadipocyte isolation and differentiation

2-4 week old C57Bl/6J (WT), Fasn^Flox/Flox^, or cAdFasnKO pups were euthanized and subcutaneous inguinal fat pads were dissected and placed in Hanks’ Balanced Salt Solution (HBSS). The fat pads were digested with 1.5mg/ml Collagenase (Sigma #C6885) with 2% BSA for 40 minutes at 37°C with shaking. The digested solution was filtered through a 100µm filter, centrifuged, and the pellet resuspended in red blood cell lysis buffer. After 2-3 minutes, the solution was centrifuged again and the pellet resuspended in growth media consisting of DMEM/F12 with 10% fetal bovine serum and 1% (v/v) penicillin/streptomycin. The cells were filtered through a 70µm filter and grown to confluency. Preadipocytes were used no later than passage 1 for these experiments. To induce differentiation (day 0), preadipocytes were grown to full confluency and cultured in induction medium containing growth media supplemented with 5□μg/mL insulin, 1□μM dexamethasone, 0.5□mM 3-isobutyl-1-methylxanthine, 60□μM indomethacin and 1□μM rosiglitazone. Forty-eight hours later the media was changed to Day 2 media consisting of growth media with 5□μg/mL insulin. The same media was replaced 48 hours later (Day 4). Adipocytes were considered fully differentiated at Day 5 and were maintained in regular growth media thereafter. Adipocytes were harvested on Day 6 or Day 7. For fatty acid (FA) supplementation experiments, a 20X stock solution containing 4mM sodium palmitate, 4mM oleic acid, and 10% fatty acid free BSA dissolved in DMEM/F12 was prepared. The FAs were conjugated to BSA by heating at 56°C for ∼1 hr and the pH was adjusted to ∼7.4. The FA mixture was added to the media at 1X concentration (200 μM sodium palmitate, 200 μM oleic acid, 0.5% BSA) on Day 0, Day 2, or Day 5 of differentiation and replaced every other day.

For ACC inhibition experiments, WT primary preadipocytes were differentiated as described above. On day 5, 10μM firsocostat (ND-630, Selleckchem) dissolved in DMSO was added for 48 hours. In experiments with co-treatment with insulin, 1μM insulin and 15μM firsocostat were added for 24, 48, or 72 hours beginning on day 5.

### Autophagy Flux Assay/LC3II Turnover Assay

#### Cultured adipocytes

For autophagy activation, cultured adipocytes were washed with PBS and incubated in HBSS for 4 hours. For autophagy inhibition, adipocytes were cultured in growth media supplemented with 50 μM chloroquine (CQ) (Sigma 6628) for 4 hours. For the combination treatment, 50uM CQ was added to HBSS and the cells were incubated for 4 hours. Control cells were incubated in growth medium. The synthesis ratio, described in (Klionsky et al., 2021) was calculated as the change in LC3II in HBSS vs. control or HBSS + CQ vs. CQ for each genotype. Similarly, the degradation ratio was calculated as the change in LC3II in the CQ vs. control or HBSS + CQ vs. HBSS condition.

#### Adipose Tissue Explants

Freshly dissected subcutaneous adipose tissue depots from individual Fasn^Flox/Flox^, or cAdFasnKO mice were washed in PBS, minced into ∼20mg pieces, and divided into two culture dishes. One dish contained DMEM/F12 and the other contained DMEM/F12 with 50uM CQ, thus each mouse/fat pad was its own control. The explants were incubated at 37°C with 5% CO_2_ for 2 hours. After 2 hours, the explants were washed briefly in PBS and protein was isolated as described.

### Isolation of adipocytes vs. stromal vascular fraction (SVF)

Subcutaneous white adipose tissue fat pads were dissected and placed in Hanks’ Balanced Salt Solution. The fat pads were digested with 1.5mg/ml Collagenase (Sigma #C6885) for 30 minutes at 37°C with shaking. The digested solution was filtered through a 100µm filter and centrifuged for 5 minutes at 500g. The resulting top layer (fat cake) was transferred to a new tube and processed for western blotting as described below. The pellet (SVF) was resuspended in protein homogenization buffer and similarly processed for western blotting.

### Electron Microscopy

Fasn^Flox/Flox^ and cAdFasnKO preadipocytes were differentiated and on day 7 were fixed by first removing half the plate media and then adding an equal volume of 2.5% glutaraldehyde (v/v) in 1 M Na phosphate buffer (pH 7.2) for 10 minutes before being transferred to pure 2.5% glutaraldehyde (v/v) in 1 M Na phosphate buffer (pH 7.2) for 1 hr. Next the cell plates were briefly rinsed (3×10mins) in 1 M Na phosphate buffer (pH 7.2) and post-fixed for 1 hr in 1% osmium tetroxide (w/v) in dH_2_O. Samples were then washed three times with dH_2_O for 10 mins and then cells were scraped off the bottom of the wells with a soft plastic spatula, collected in a microfuge tube, and pelleted by centrifugation. Samples were washed three times with dH_2_O for 10 minutes and dehydrated through a graded series of ethanol (10, 30, 50, 70, 85, 95% for 20 min each) to three changes of 100% ethanol. Samples were infiltrated first with two changes of 100% Propylene Oxide and then with a 50%/50% propylene oxide/SPI-Pon 812 resin mixture overnight. The following morning the cell pellets were transferred through four changes of fresh SPI-pon 812-Araldite epoxy resin and finally embedded in tubes filled with the same resin and polymerized for 48 hours at 70°C. The epoxy blocks were then trimmed, and ultrathin sections were cut on a Reichart-Jung ultramicrotome using a diamond knife. The sections were collected and mounted on copper support grids and contrasted with lead citrate and uranyl acetate. The samples were examined on a Philips CM 10 using 100 Kv accelerating voltage. Images were captured using a Gatan TEM CCD camera.

### Real-time quantitative PCR

RNA was isolated from cultured adipocytes or tissue with Trizol, following the manufacturer’s instructions. cDNA was synthesized from 1µg RNA using Bio-Rad iScript cDNA kit. qPCR was performed using Bio-Rad iTaq SYBR Green Supermix on a BioRad CFX97 thermocycler and analyzed using the ΔΔCt method. For cultured adipocytes, 18s was used for normalized, and for tissues, the average of *18s* and *B2m* was used for normalization. Primer sequences are listed in Table

### Western blotting

Cultured adipocytes were harvested in 50 mM Tris-HCl, pH 7.6, 150 mM NaCl, 5 mM EDTA, 1% Triton X-100 supplemented with Halt protease and phosphatase inhibitors (ThermoScientific). The homogenate was spun at 6,000g for 15 minutes at 4°C. The resulting supernatant was considered the “Triton X-100 soluble fraction,” and the pellet considered the “Triton X-100 insoluble fraction.” To solubilize the pellet, 2.5X Laemmli buffer with β-mercaptoethanol was added. For adipose tissue, fat pads or explants were suspended in 50 mM Tris-HCl, pH 7.6, 150 mM NaCl, 5 mM EDTA with Halt protease and phosphatase inhibitors and homogenized with the Qiagen TissueLyser. Homogenates were spun at 6,000g for 15 minutes at 4°C and the resulting infranatant, between the pellet and overlying fat cake was collected. Protein concentrations were determined by BCA (Pierce). Proteins were prepared for electrophoresis by adding Laemmli buffer with β-mercaptoethanol and heating to ∼90°C for 10 minutes. Proteins were resolved by SDS-PAGE and blotted with the following antibodies: anti-Fasn (CST #3180); anti-Ucp1 (Abcam #10983); anti-p62 (CST #23214); anti-alpha-tubulin (Sigma #T5168); anti-LC3B (CST #83506), anti-LC3A/B (CST #12741); anti-Gapdh (CST #8884); anti-malonylated lysines (PTM Biolabs #901); anti-vinculin (CST #18799); anti-Gabarap (CST #13733)

### Histology

Freshly dissected adipose tissue pieces were fixed in 4% paraformaldehyde overnight and embedded in paraffin. Sections were stained with H&E and anti-p62 (CST #23214) at the UMass Medical School Morphology Core. Images were taken with a Leica DM2500 LED microscope equipped with a Leica MC170 HD camera.

### Immunofluorescence

Preadipocytes were grown on glass coverslips and differentiated as described. On day 6 or 7, differentiated adipocytes were fixed in 4% paraformaldehyde for 10 minutes at room temperature, washed, and permeabilized with ice-cold 100% methanol for 10 minutes. The cells were incubated with blocking solution containing 2% normal goal serum, 1% BSA, 0.1% Triton X-100, and 0.05% Tween 20. Primary antibodies were diluted in blocking solution and incubated overnight at 4°C. Secondary antibodies were diluted in blocking solution and incubated at room temperature for 1 hour. Nuclei were stained with 1ug/ml DAPI (Invitrogen), and coverslips were mounted on slides with Prolong Glass mountant (Invitrogen).

Images were taken with a Leica TCS SP8 confocal microscope. Colocalization analysis was performed with the JACoP plugin in ImageJ. Antibodies: rabbit anti-p62 (CST #23214); mouse anti-p62 (R&D #MAB8028); rabbit anti-LC3A/B (CST #12741); mouse anti-LC3B (CST #83506); Alexa Fluor-594 goat anti-rabbit and Alexa Fluor-488 goat anti-mouse (Invitrogen)

### Lipidomics

Lipid species were analyzed using multidimensional mass spectrometry-based shotgun lipidomic analysis (Han, 2016). In brief, homogenate of adipocytic sample containing 0.2□mg of protein as determined by the Pierce BCA assay was accurately transferred to a disposable glass culture test tube. A pre-mixture of lipid internal standards (IS) was added prior to conducting lipid extraction for quantification of the targeted lipid species. Lipid extraction was performed using a modified Bligh and Dyer procedure (Wang and Han, 2014), and each lipid extract was reconstituted in chloroform:methanol (1:1, v:v) at a volume of 500 µL/mg protein.

For shotgun lipidomics, individual lipid extract was further diluted to a final concentration of ∼500 fmol total lipids per µL. Mass spectrometric analysis was performed on a triple quadrupole mass spectrometer (TSQ Altis, Thermo Fisher Scientific, San Jose, CA) and a Q Exactive mass spectrometer (Thermo Scientific, San Jose, CA), both of which were equipped with an automated nanospray device (TriVersa NanoMate, Advion Bioscience Ltd., Ithaca, NY) as described (Han et al., 2008). Identification and quantification of lipid species were performed using an automated software program (Wang et al., 2016; Yang et al., 2009). Data processing (e.g., ion peak selection, baseline correction, data transfer, peak intensity comparison and quantitation) was performed as described (Yang et al., 2009). The results were normalized to the protein content (nmol or pmol lipid/mg protein).

### Acetyl-CoA and malonyl-CoA measurements

Malonyl- and acetyl-CoA were extracted with 0.3 M perchloric acid and analyzed by LC-MS/MS using a method based on a previously published report by (Gao et al., 2007). The extracts were spiked with ^13^C_2_-Acetyl-CoA (Sigma, MO, USA), centrifuged, and filtered through the Millipore Ultrafree-MC 0.1 µm centrifugal filters before being injected onto the Chromolith FastGradient RP-18e HPLC column, 50 × 2 mm (EMD Millipore) and analyzed on a Waters Xevo TQ-S triple quadrupole mass spectrometer coupled to a Waters Acquity UPLC system (Waters, Milford, MA).

### Primer Sequences

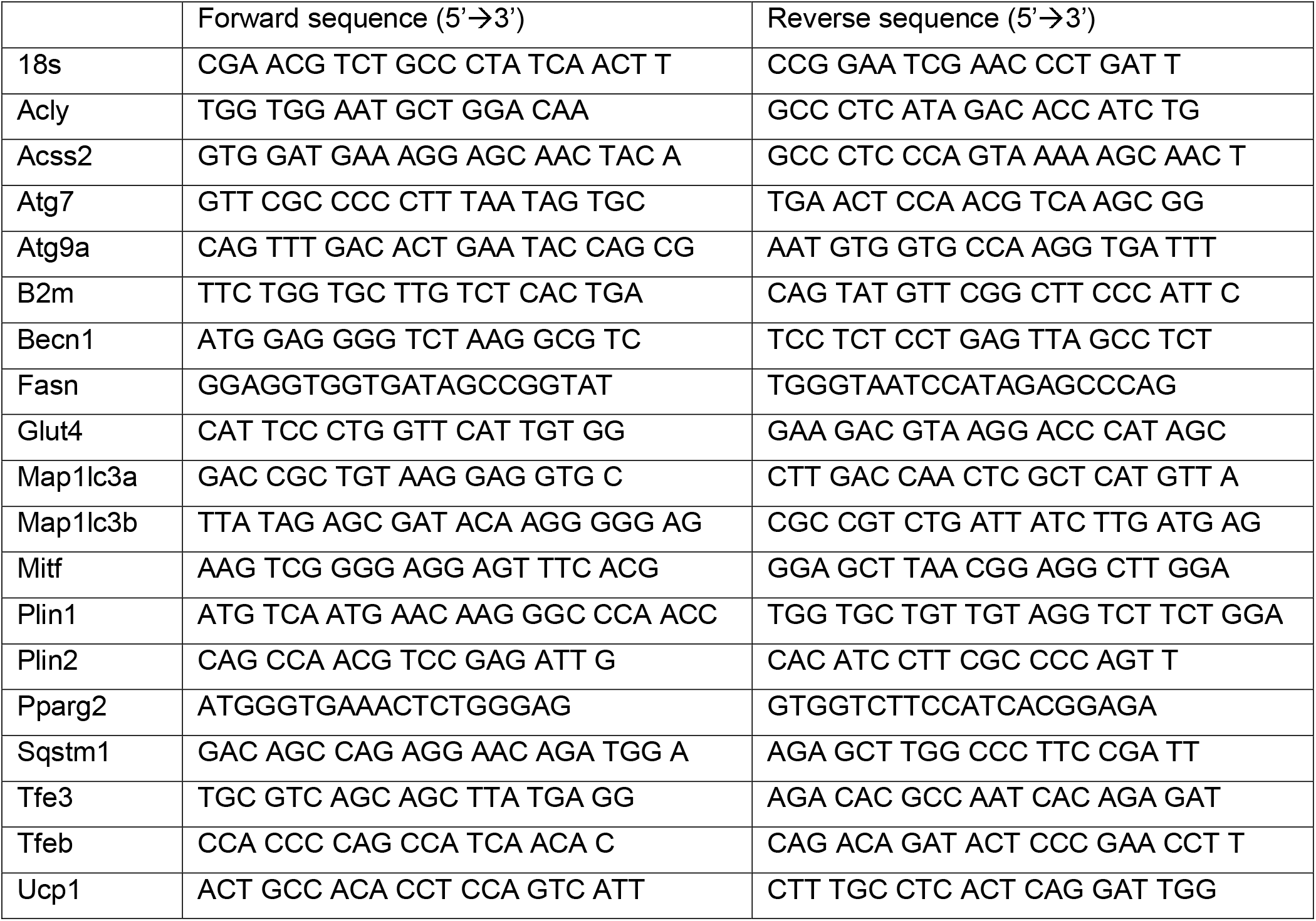

## Supplemental Figure Legends

1. A) Immunofluorescence of Fasn^Fl/Fl^ and cAdFasnKO adipocytes labeled with LC3 or p62. Scale bar = 50um. B) Related to Figure 1G. Quantification of LC3II levels normalized to alpha-tubulin. 2-way ANOVA with Tukey post hoc: *<0.05, **<0.01, ****<0.0001

2. Related to Figure 2H. Quantification of LC3II (A) and p62 (B) levels normalized to α-tubulin. 2-way ANOVA with Tukey post hoc: *<0.05, **<0.01, ****<0.0001. All data are means +/- SE.

3. A) Light microscopy of Fasn^Fl/Fl^ and cAdFasnKO adipocytes supplemented with 200uM each of palmitate and oleate beginning 48 hours after the onset of differentiation (day 2). B) Western blots of Triton X-100-soluble and -insoluble protein fractions. C) Quantification of soluble p62 and insoluble p62 from blot in B. D) Quantification of LC3 from blot in B. Data are means +/- SE. Two-way ANOVA with Tukey post hoc: **<0.01, ***<0.001, ****<0.0001.

4. Related to Figure 5. A) Double bond composition of triglycerides and B) phophatidylethanolamine (PE). Top panels are in vitro cultured adipocytes and bottom panels are from in vivo adipose tissue. Data are means +/- SE. t tests: *<0.05, ***<0.001, ****<0.0001. C) Heat map displaying fold changes of the percent of total phosphatidylinositol species from Fasn^Fl/Fl^ vs. cAdFasnKO <u>adipocytes</u> and iAdFasnKO <u>iWAT</u>. t tests: * <0.05.

## Notes

### Competing Interest Statement

The authors have declared no competing interest.

