## Supplementary figures and images for "De novo lipogenesis fuels adipocyte autophagosome membrane dynamics"

### Supplemental Figures

#### Supplemental Figure 1.

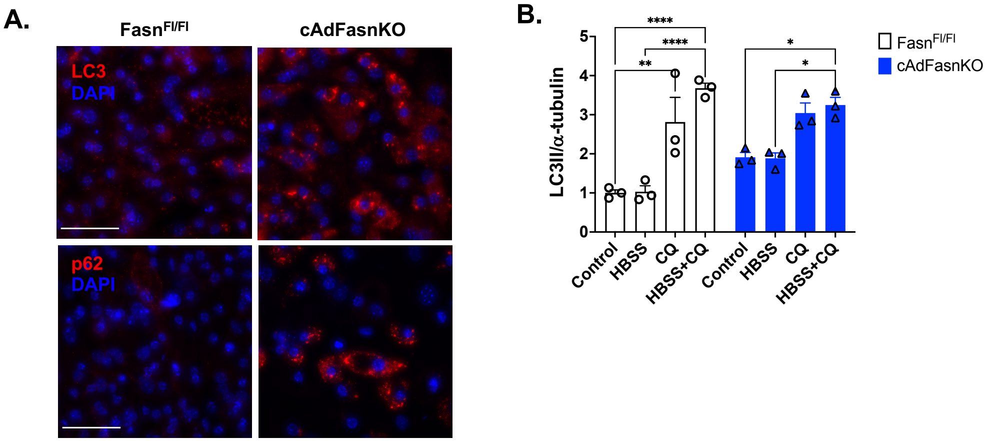

##### Supplemental Figure 2.

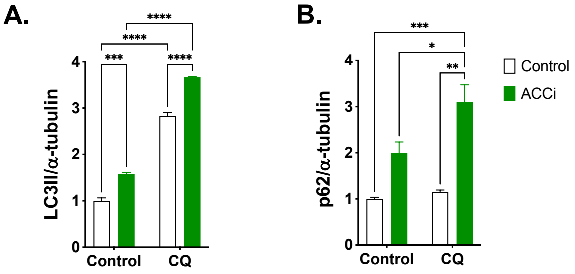

##### Supplemental Figure 3.

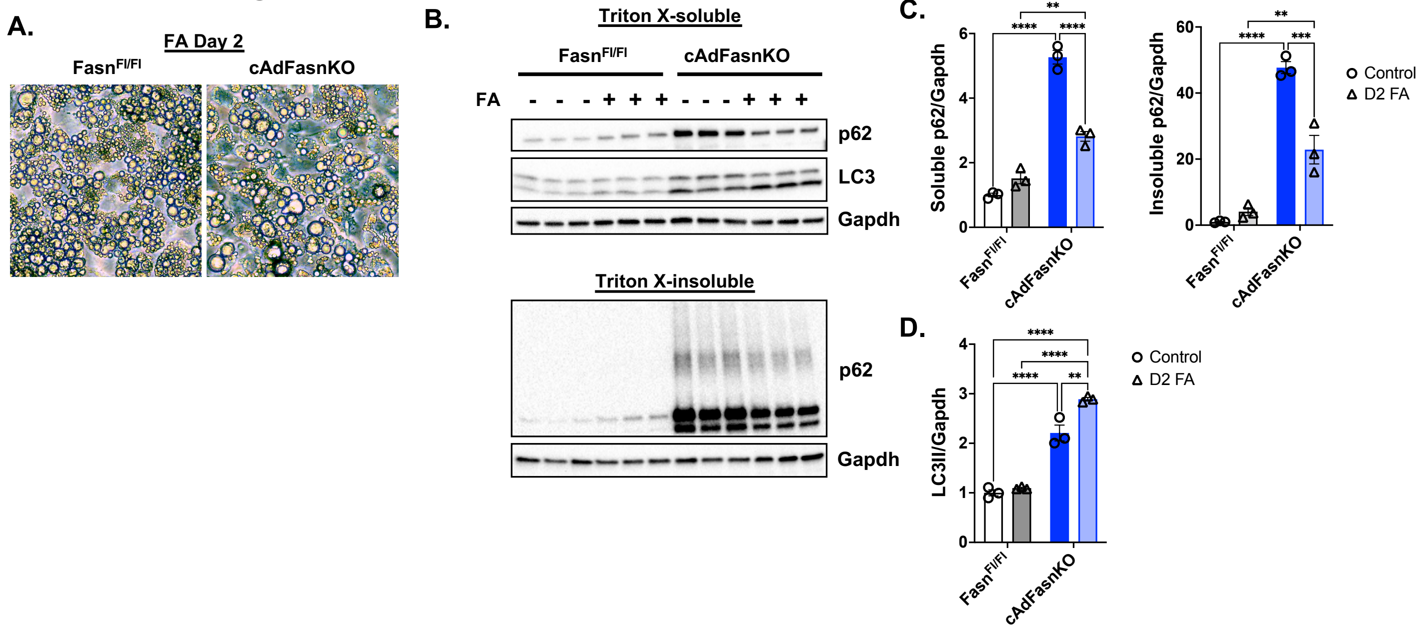

#### Supplemental Figure 4

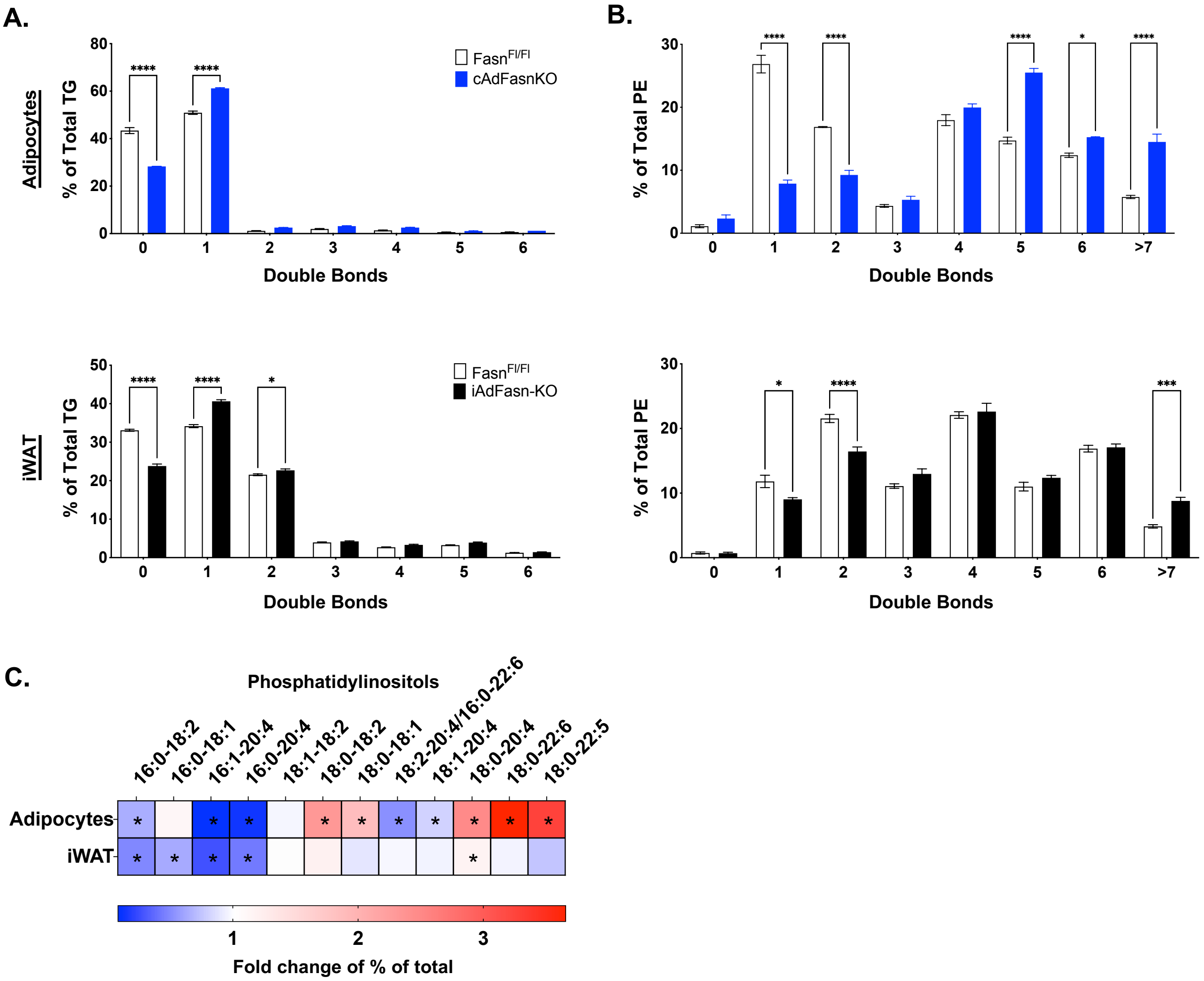
